# Prenatal stress increases corticosterone levels in offspring by impairing placental glucocorticoid barrier function

**DOI:** 10.1101/2024.10.31.621366

**Authors:** Can Liu, Hongya Liu, Hongyu Li, Deguang Yang, Ye Li, Rui Wang, Jiashu Zhu, Shuqin Ma, Suzhen Guan

**Author notes:** Corresponding authors at School of Public Health of Ningxia Medical University, Yinchuan, 750004, Ningxia, China. E-mail addresses (S. Guan). These authors have contributed equally to this work (Can Liu, Hongya Liu).

## Abstract

To investigate the association between prenatal stress (PS) and corticosterone, and its influence on DNA methylation of genes related to the placental glucocorticoid (GC) barrier, including 11β-HSD2, P-gp, NR3C1, and FKBP5. The PS model was established through chronic unpredictable mild stress (CUMS). DNA methylation of GC-related genes was analyzed using reduced representation bisulfite sequencing (RRBS), and the results were confirmed using MethylTarget™ sequencing. The mRNA and protein expression levels of these genes were detected through qRT-PCR and Western blotting, respectively. Plasma corticosterone levels are elevated in pregnant female rats exposed to PS conditions and their offspring. Compared to the offspring of the prenatal control (OPC) group, the offspring of the prenatal stress (OPS) group showed down-regulation in both mRNA and protein expression of DNMT 3A and DNMT 3B, while up-regulation was observed in the expression of DNMT1. RRBS analyses identified P-gp and FKBP5 as hypermethylated genes, including a total of 43 differentially methylated sites (DMS) and 2 differentially methylated regions (DMR). MethylTarget™ sequencing revealed that both genes had 15 differentially methylated CpG sites. This study provides preliminary evidence that PS disrupts the placental GC barrier through abnormal gene expression caused by hypermethylation of GC-related genes, resulting in elevated corticosterone levels in offspring and affecting their growth and development.

## 1. Introduction

Pregnancy is a physiological process widely regarded as “natural” and experienced by women throughout their lives, yet it is also considered a persistent source of stress. Pregnant women may encounter various stressors, including environmental toxins, family conflicts, work-related pressures, and mental trauma, which in excess can disrupt their psychological well-being. Numerous epidemiological and case-control studies have demonstrated that maternal psychological and social stress during pregnancy (prenatal stress, PS) not only increases health risks for mothers [1,2], but also elevates the likelihood of negative pregnancy outcomes such as miscarriage, preterm birth, and low birth weight due to prenatal stress[3–6]. As previously mentioned, anxiety caused by prenatal stress has been recognized as a significant risk factor for abnormal health conditions in offspring including emotional disorders and depressive behaviors[7].

Barker’s Developmental Origins of Health and Disease (DOHaD) hypothesis[8,9] suggests that PS can lead to cognitive impairments and mental disorders in children by altering physiological and metabolic responses. Excessive PS is associated with structural and physiological changes in offspring, including epigenetic modifications[10–13].

Currently, significant attention has been focused on the association between stress exposure during pregnancy and adverse health outcomes; however, the underlying mechanisms of PS transmission remain unclear. Cortisol, a well-known stress hormone, primarily functions as a glucocorticoid (GC) in mammals. The placenta acts as a barrier for protecting the developing fetus from the adverse effects of excessive maternal glucocorticoids. One widely accepted mechanism suggests that maternal cortisol disrupts the placental GC barrier under high-stress conditions and exposes the fetus to cortisol through placental blood flow[14]. In a state of PS, persistent activation of the hypothalamic-pituitary-adrenal (HPA) axis can lead to excessive secretion of GCs and subsequent dysfunction in various systems such as nervous, endocrine, and immune systems[15]. Notably, impaired integrity of the placental GC barrier has been associated with intrauterine growth retardation and increased risk for chronic diseases later in life[16].

Vuppaladhadiam et al. found a link between plasma and placental cortisol in prenatal stress (PS), hinting at epigenetic impacts on child development and health[17]. Key regulators of maternal and fetal cortisol levels include 11β-hydroxysteroid dehydrogenase 2 (11β-HSD2)[18] and p-glycoprotein (P-gp). 11β-HSD2 inactivates cortisol, while P-gp transports it back to the maternal circulation[19,20]. Chronic stress may alter glucocorticoid sensitivity through DNA methylation changes in stress-related HPA axis genes[21].

Prenatal stress (PS) is known to alter glucocorticoid receptor (GR) gene expression, including NR3C1 and FKBP5, crucial for HPA axis regulation and stress response[22]. Changes in NR3C1 methylation due to stress can affect neonatal self-regulation[23,24] and are linked to maternal depression and adversity[25,26]. Despite the known connection between maternal care and NR3C1 methylation in offspring, the full impact of epigenetic changes on placental GC barrier genes due to PS requires further study.

Our study used a chronic unpredictable mild stress (CUMS) model to investigate the effects of PS on pregnant rats. We examined the DNA methylation of four placental GC barrier-related genes (11β-HSD2, P-gp, NR3C1, and FKBP5) and their association with HPA axis activity. Based on abnormal placental DNA methylation observed in our study, we hypothesized that PS would lead to elevated corticosterone levels associated with these genes, thereby elucidating the underlying mechanism linking maternal/fetal cortisol elevation with pregnancy-related stress.

## 2 Materials and methods

### 2.1 Chemicals and reagents

The Iodine-131 cortisol radioimmunoassay kit was obtained from the Beijing North Institute of Biotechnology. The RNA extraction kit, first strand cDNA synthesis kit, and real-time PCR reaction kit were purchased from Beijing Tiangen Biochemical Technology Co., Ltd. RIPA lysis solution was obtained from Nanjing KGI Biotechnology Co., Ltd. The whole protein extraction kit and bicinchoninic acid (BCA) protein content detection kit were utilized. Sodium dodecyl sulfate-polyacrylamide gel electrophoresis (SDS-PAGE) gel preparation kit and substrate chemiluminescence (electro-chemo-luminescence, ECL) kit were purchased from Nanjing KGI Biotechnology Co., Ltd.

Primary antibodies including anti-β-actin (AF7018), anti-DNA methyltransferases (DNMT) 3A (DF7226), anti-DNMT 3B (AF5493), anti-DNMT1 (DF7376) and Goat-Rabbit IgG (S0001) were purchased from Affinity Technology Co., Ltd. Anti-11β-HSD2 antibody was acquired from Novus Technology Co., Ltd. (NBP1-39478). The anti-P-gp (ab170904) and anti-NR3C1 (ab183127) were purchased from Abcam Technology Co., Ltd. The anti-FKBP5 (00083356) was purchased from Proteintech Technology Co., Ltd.

### 2.2 CUMS Procedure

Sixteen healthy adult female Sprague-Dawley (SD) rats weighing (200 ± 20) g and twelve male Sprague-Dawley (SD) rats weighing (220 ±20) g were obtained from the Experimental Animal Center of Ningxia Medical University, with an experimental animal certificate number SYXK(Ning)2020-0001. All procedures were approved by the Animal Care and Use Committee of Ningxia Medical University and conducted by the National Institutes of Health Guide for the Care and Use of Laboratory Animals (IACUC-NYLAC-2020068). Rats were group-housed in cages under controlled temperature conditions (22∼24°C), with a 12-hour light/dark cycle.

Female rats were randomly assigned to either a prenatal stress (PS) group or a prenatal control (PC) group, each consisting of eight rats. Male rats were randomly divided into a control mating group (four rats) or a stress mating group (eight rats). Mating for the PS group commenced on day 7 after exposure to chronic unpredictable mild stress. Before gestation, female rats from both groups mated with males from their respective mating groups. Pregnancy was determined by a daily vaginal smear check of the pubic plug, and the birth date of the offspring was designated as gestational day 0 (GD 0), subsequently, male and female rats were separated. After gestation, the rats were returned to their original breeding environment. During mating, the PS group rats experienced continuous stress stimulation. To induce chronic stress conditions, we employed the chronic unpredictable mild stress (CUMS) procedure based on a previously described method with minor modifications and additions[2,27]. The PS group rats were exposed to one randomly selected stressor out of seven different types daily for four weeks. These stressors included crowded environments for 24 hours, physical restraint (2 hours), filthy cages for 24 hours (60-70% humidity, 12 hours), hot water swimming at 45°C for 5 minutes, rocking cages for 30 minutes, ice bath at 4°C for 5 minutes, and tail squeezing for 2 minutes. Each day, one of these seven different stressors was randomly administered. The duration of the daily exposure to the twenty-four-hour stressors was from morning at 10:00 until the next morning at 10:00; all other stressors occurred between morning at 10:00 and noon. Throughout this process, stressed rats were temporarily relocated to another room with identical light intensity and temperature before being returned to the main room following stimulation.

### 2.3 Placental sampling

After 18 days of gestation, six female rats (randomly chosen three rats from both the PS group and PC group) were anesthetized with isoflurane, and the uterus tissues were promptly removed. The uterus-embryo mixture was pre-cooled in PBS solution, followed by removal of the uterus and fetal membrane tissues, and subsequent separation of the placenta. Three placental tissues from each group were immediately frozen in liquid nitrogen for storage at -80°C to facilitate subsequent examination.

### 2.4 Allocation of the offspring into groups

On postnatal day 0 (PND 0), the day of birth, dams, and pups remained together until weaning on PND 21, when male and female pups were separated. Offspring selected from the PC group and PS group were randomly assigned to the offspring of the prenatal control (OPC) group (n = 10; 5 males vs 5 females) or the offspring of the prenatal stress (OPS) group (n = 10; 5 males vs 5 females). All experimental offspring were housed together in the same room with ad libitum access to water and food. Body weight measurements were taken, and blood samples were collected from the inner canthal vein on PND28 and PND42. A schematic diagram of the experimental procedure is shown in Fig.S1.

### 2.5 Measurement of selected indicators

#### 2.5.1 Measurement of plasma corticosterone concentration

The successful establishment of the CUMS model was confirmed by measuring the plasma corticosterone concentration in female rats at different time points during stress. Blood samples (1 mL) were collected from the inner canthus vein of rats on the day before the first stress (baseline), as well as on days 1, 7, 14, 21, and 28 during the stress period. The plasma was separated and stored at -80°C and corticosterone levels were measured using a ^131^I cortisol radioimmunoassay (RIA) kit according to the manufacturer’s instructions. The intra-assay variability of RIA ranged from 3.2% to 4.7%. Plasma corticosterone concentrations were calculated using a conversion formula: corticosterone concentration = cortisol concentration ×50[28].

#### 2.5.2 Reduced representation bisulfite sequencing

Genomic DNA from placental samples of two offspring groups (three placental samples in the OPC group and three placental samplings in the OPS group) was extracted using a magnetic universal genomic DNA Kit following the manufacturer’s protocol. RRBS library preparation was performed using Acegen Rapid RRBS Library Prep Kit according to the manufacturer’s instructions.

#### 2.5.3 Illumina Hiseq sequencing platform

RRBS sequencing was conducted at Biomarker Technologies Co.,LTD. RRBS sequencing was conducted at biomarker technologies where genomic DNA (100 ng) was digested with MspI, end-repaired, tailed with dA nucleotides, and ligated to adapters modified with methylcytosine residues. Genomic DNA purification was carried out using Agencourt® AMPure® XP Nucleic Acid Purification Kit followed by bisulfite treatment utilizing ZYMO EZ DNA Methylation-Gold Kit. Illumina double index primers with an eight-base pair sequence were used for PCR amplification consisting of twelve cycles. The quality analysis of libraries involved an Agilent 2100 bioanalyzer (Agilent Technologies, Santa Clara, CA, USA) while quantification of DNA fragments ranging from sizes between25 bp to12,000 bp utilized ABI StepOnePlus real-time PCR system (Thermo Fisher Scientific). Fragments were sequenced using a 150×2 paired terminal sequencing scheme on the Illumina Novase Q6000 analyzer (Illumina HiSeq sequencing platform) to obtain the sequencing results. Subsequently, methylation analysis was performed on the promoter and CpG island region by detecting the proportion of CG, CHG, and CHH in the methylated C base, the average methylation level of the C base, and the distribution of different types of methylation. Regional methylation profiles were calculated based on methylation levels and CpG density for specific regions, with CpG density defined for each CpG site within a 200 bp window. Differential Methylation Region (DMR) analysis was conducted using a sliding window method to calculate the methylation difference between placental samples from OPC and OPS groups.

#### 2.5.4 Validation of DNA methylation status and detection of the expression levels of the identified markers

The DNA methylation status of the P-gp (abcb1a) and FKBP5 intron was analyzed using the MethylTarget™ assay (targeted bisulfite sequencing). Target genes with the greatest differences in DNA methylation were subjected to further analysis (Table S1). Six placentas in the OPC group and six placentas in the OPS group) was used to validate the results of RRBS and to detect the expression of these differentially methylated genes. Sample acquisition, DNA extraction, and preservation were performed as mentioned above.

Firstly, 40 ng of genomic DNA underwent bisulfite conversion using the EZ DNA Methylation-Gold™ kit (Zymo Research) according to the manufacturer’s instructions. PCR amplification was then performed to amplify targeted DNA sequences. The resulting products were sequenced on an Illumina MiSeq benchtop sequencer. Primer sequences for target regions within FKBP5 and P-gp (abcb1a) loci are listed in Table 1 below. FastQC software was used for quality control assessment of sequencing reads. The Filtered reads were subjected to read recalibration with USEARCH and then mapped to the genome by Blast.

**Table 1.**
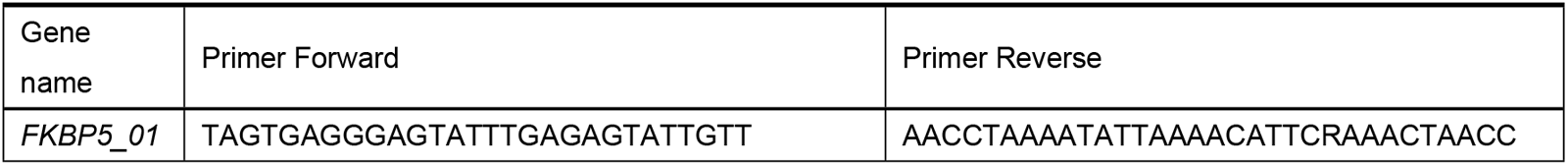
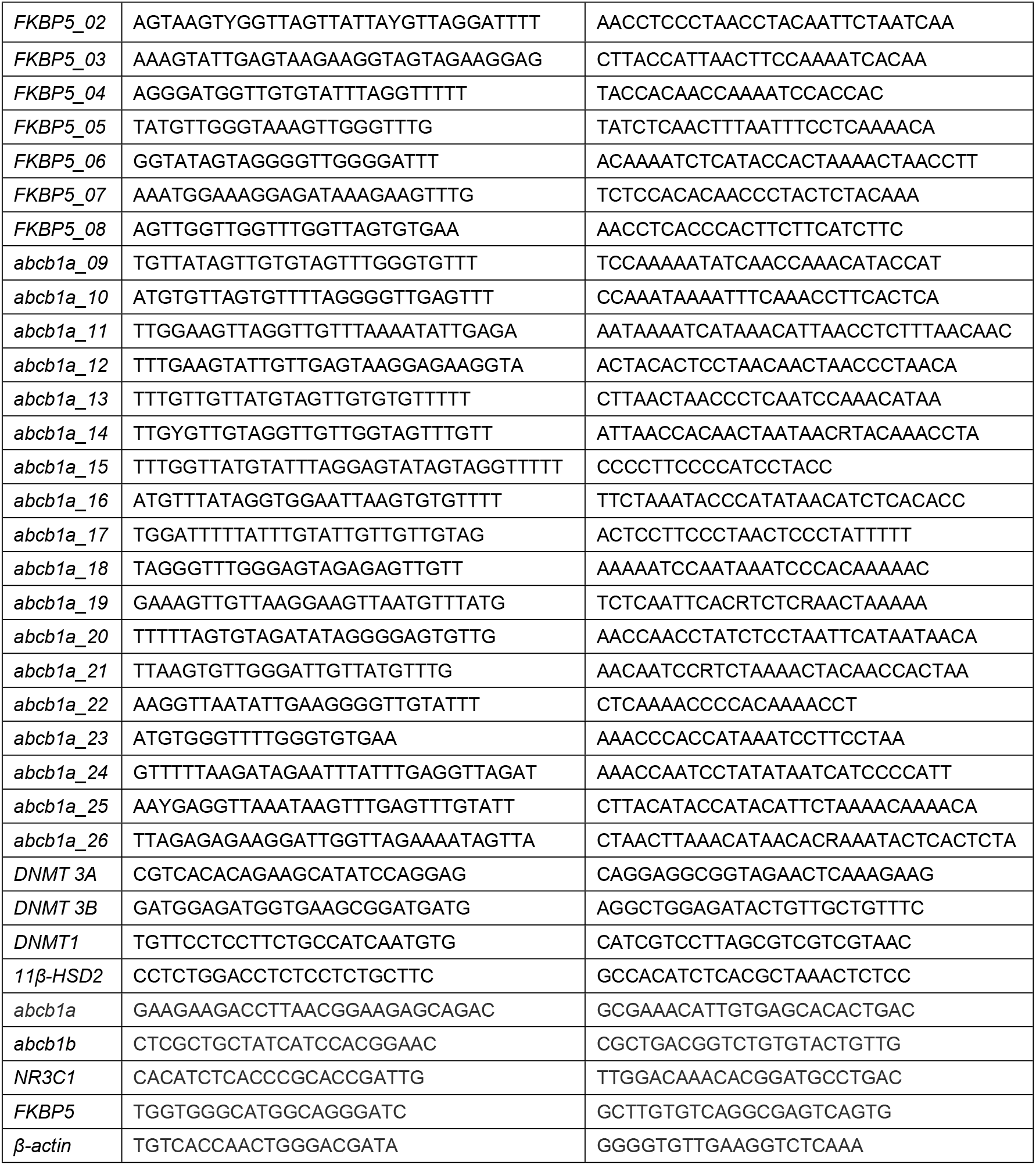
P Primers in MethylTarget sequencing for validation of DNA methylation status and in qRT-PCR for detection of gene transcription.

#### 2.5.5 Quantitative real-time PCR (qRT-PCR)

The RNA was extracted from isolated placenta tissues following the protocol of the total RNA extraction kit. The extracted RNA was quantified using a nucleic acid protein quantitative instrument, and samples with an A260/280 ratio between 1.8 and 2.0 were selected for further analysis. A two-step Real-time Quantitative PCR (RT-qPCR) assay was performed to analyze gene expression levels related to DNA methyltransferases (DNMT)3A, DNMT3B, DNMT1, 11β-HSD2, abcb1a, abcb1b, NR3C1, and FKBP5. The cDNA was generated from total RNA derived from placental tissue (1000ng) in a total volume of 20 µL using a cDNA synthesis kit. PCR reactions were carried out using primers (listed in Table 1) and SuperReal PreMix Plus(SYBR Green) reagent under specific conditions: the pre-denaturation step lasted 15 minutes at 95°C, followed by the denaturation step lasting 10 seconds at 95 °C, the annealing step lasting 20 seconds at 55 °C, and extension step lasting 30 seconds at 70 °C, for 40 cycles. Finally, the relative expression level of the target gene was calculated using the 2^-ΔΔCt^ method.

#### 2.5.6 Western Blot

The placenta tissue was subjected to protein extraction using a cold Radioimmunoprecipitation assay (RIPA) lysis buffer system. The protein concentration of the samples was determined using a BCA assay kit. Equal amounts of protein were separated by SDS-PAGE at a 10% concentration and subsequently transferred onto a PVDF microporous membrane. Following blocking with 5% skim milk, the membranes were incubated overnight at 4°C with antibodies of appropriate concentrations, including anti-DNMT1 (1:1000), anti-DNMT3A (1:1000), anti-DNMT3B (1:1000), anti-11β-HSD2 (1:2000), anti-P-gp (1:2000), anti-NR3C1 (1:2000), anti-FKBP5 (1:2000), or anti-β-actin Protein antibody (1:2000). Afterward, secondary antibodies were applied at room temperature for 1 hour. Visualization of the blots was achieved using ECL Plus detection reagent from Jiangsu Kaiji Biotechnology Co., LTD.

### 2.6 Statistical analysis

Statistical analysis was performed on all data using the Statistical Package for the Social Sciences (SPSS) version 26.0, while graph construction utilized GraphPad Prism version 8.0. Differences in plasma corticosterone levels during prenatal stages were analyzed through repeated measurements analysis of variance and Student’s t-test for multiple comparisons at different time points. RT-PCR and Western blotting data were analyzed using Student’s t-test to compare the two offspring groups.

## 3 Results

### 3.1 CUMS increases prenatal plasma corticosterone levels

Repeated-measures ANOVA results showed that chronic stress had a significant effect on prenatal corticosterone concentration (*F*=7.632, *P*=0.020), and the concentration of corticosterone in the PS group changed with the increase of stress duration ((*F*=6.871, *P*=0.001). In addition, there was a significant interaction between stressors and time (*F*=3.141, *P*=0.015). The plasma corticosteroid concentrations in the PS group were higher than that in the PC group on days 14, 21, and 28 of stress (*t*_*14*_=-2.811, *P*=0.018; *t*_*21*_=-3.398, *P*=0.007; *t*_*28*_=-3.060, *P*=0.012). The results suggest that the PS group was stressed during pregnancy. Plasma corticosterone concentrations were not statistically significant between the PC and PS groups at baseline examination *(t=-1*.*607, P=0*.*139)* and on day one after exposure to stress *(t=-0*.*359, P=0*.*727)* (Fig.S2).

### 3.2 Effects of PS on offspring body weight and plasma corticosterone

Repeated measures analysis of variance showed significant differences in corticosterone concentrations between the OPS and the OPC group (*F*=20.588; *P*=0.000). Corticosteroid concentrations in the OPS group changed with increasing birth time (*F*=8.783; *P*=0.008). No significant difference was found in the interaction between the treatment groups times (*F*=0.903; *P*=0.355). The plasma corticosterone concentrations in the OPS group were higher than that in the OPC group at PND28 (*t*=-2.537; *P*=0.027, Fig.1A) and PND42 (*t*=-3.764; *P*=0.003, Fig.1B). The results suggest that the OPS group were in a state of high plasma corticosterone level. Repeated measures ANOVA of the two groups showed a significant effect on offspring body weight (*F*=14.384, *P*=0.001), and there was a significant difference between time and body weight, but not between interaction and body weight (Time: *F*=641.390, *P*=0.000; Interaction: *F*= 0.750, *P*=0.398). The body weight in the OPS group was lower than in the OPC group at PND28(*t*=3.645; *P*=0.002, Fig.1C) and PND42 (*t*=3.044; *P*=0.008, Fig.1D).

**Fig.1.**
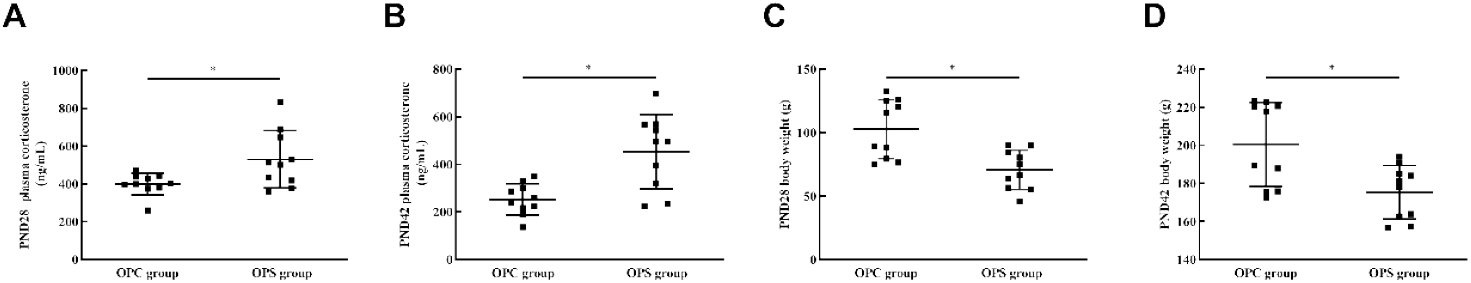
Effects of PS on offspring’s body weight and plasma corticosterone. The figure showed that offspring after exposure to PS were measured plasma corticosterone and body weight on PND28 and PND42. (A, B) Plasma corticosterone concentrations in offspring exposed to prenatal stress were higher on PND28 and PND42. (C, D) Body weight in offspring exposed to prenatal stress was lower on PND28 and PND42. Data are represented as mean ± SD (N=10 per group). *Statistically significant difference compared with the OPC group (*P*<0.05).

### 3.3 The effects of offspring after prenatal stress during pregnancy on enzymes of DNA methylation (DNMTs)

As shown in Fig.2, the mRNA expression of *DNMT 3A, DNMT 3B*, and *DNMT1* was different between the two offspring groups, as shown in Fig.2A∼C (all *P*<0.05). Compared with the OPC group, the mRNA levels of *DNMT 3A* (*t*=3.997, *P*=0.003, Fig.2B) and *DNMT 3B* (*t*=3.210, *P*=0.009, Fig.2C) were markedly down-regulated and the mRNA expression of *DNMT1* (*t*=-2.401, *P*=0.037, Fig.2A) was up-regulated in the PS group. The protein levels of DNMT 3A (*t*=8.443, *P*=0.001, Fig.2D, F) and DNMT 3B (*t*=5.610, *P*=0.005, Fig.2D, G) were down-regulated in the OPS group, while the protein levels of DNMT1 were up-regulated (*t*=-3.601, *P*=0.023, Fig.2D, E). Overall, it is shown that PS induces changes in placental DNMTs in the offspring.

**Fig.2.**
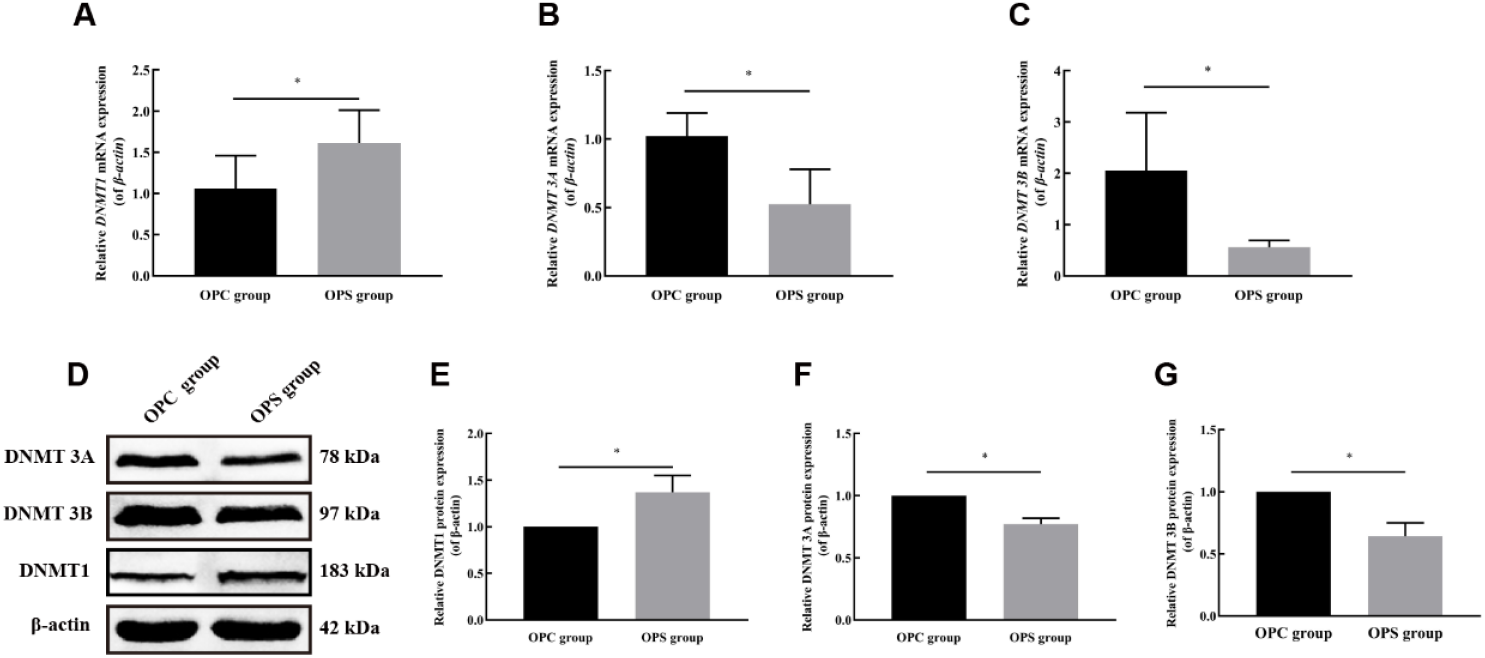
Effects of PS on offspring DNMTs in the placenta. The figure showed that the expressions of DNMT1, DNMT 3A, DNMT 3B, and β-actin were detected by (A-C) q-PCR and (D) western blot. (E-G) The levels of DNMT 3A and DNMT 3B were reduced but DNMT1 expression was increased by PS on offspring. Three biological replicates were performed. All data are shown as mean ± SD. Number of animals in each group = 3. *Statistically significant difference compared with the OPC group (P<0.05).

### 3.4 DNA methylation status in Placenta

RRBS was performed using placental tissues to identify differentially methylated CpG sites and regions. After quality control and data analysis, 222,119 differentially methylated sites (DMS) and 6905 DMRs were identified between the two groups. Compared with the OPC group, 4055 DMRs were highly methylated the in OPS group, and 2850 were low methylation. The analysis revealed that out of the total sites, 110752 were annotated with gene names, while 59778 exhibited hypermethylation and 50974 displayed hypomethylation. Table 2 shows DMR and gene numbers in differentially methylated genes between the OPC and OPS groups.

**Table 2.** Statistics of DMR in all differentially methylated gene.

| Different methylation region | Gene numbers |
| --- | --- |
| Promoter | 261 |
| 5'UTR | 60 |
| Exon | 1219 |
| 3'UTR | 121 |
| intron | 1648 |
| Distal Intergenic | 3509 |
| Downstream | 87 |
| Total | 6905 |

From all the genes with identified DMRs, there were two genes associated with placental GC barrier function: P-gp (abcb1a) and FKBP5. P-gp(abcb1a) has a total of 32 DMS Compared to the OPC group, 8 sites were hypermethylated and 5 sites were hypomethylated in the exon, and 12 sites were hypermethylated and 7 sites were hypomethylated in an intron in the OPS group. The DMS formed two DMRs, located at exon and intron regions, both with high methylation levels. FKBP5 has a total of 11 DMS (all *P*<0.05), 5 sites were hypermethylated 3 sites were hypomethylated in the intron, and 1 site was hypermethylated upstream and 2 hypermethylated downstream. Table3.

**Table 3.** Statistics of DMS and DMR in placental GC barrier-related genes FKBP5 and abcb1a.

| Gene | Position | Meth_direction | P value | Annotation |
| --- | --- | --- | --- | --- |
| *FKBP5* | 7975113 | strongHyper | 0.000388 | downstream |
| *FKBP5* | 7975116 | strongHyper | 0.00181 | downstream |
| *FKBP5* | 7977241 | strongHypo | 0.00809 | intron |
| *FKBP5* | 7977273 | strongHyper | 0.00441 | intron |
| *FKBP5* | 7978647 | strongHypo | 0.000072 | intron |
| *FKBP5* | 7991860 | strongHypo | 0.000641 | intron |
| *FKBP5* | 7991871 | strongHypo | 0.00288 | intron |
| *FKBP5* | 7992251 | strongHyper | 0.00125 | intron |
| *FKBP5* | 8000100 | strongHypo | 0.000573 | intron |
| *FKBP5* | 8002228 | strongHyper | 0.000295 | intron |
| *FKBP5* | 8019250 | strongHyper | 0.00239 | upstream |
| *abcb1a* | 22140561 | strongHyper | 0.00356 | intron |
| *abcb1a* | 22140572 | strongHyper | 0.00524 | intron |
| *abcb1a* | 22141030 | strongHyper | 0.00142 | exon |
| *abcb1a* | 22141031 | strongHyper | 0.000573 | exon |
| *abcb1a* | 22141043 | strongHyper | 0.0057 | exon |
| *abcb1a* | 22141058 | strongHyper | 0.000483 | exon |
| *abcb1a* | 22141069 | strongHyper | 0.000151 | exon |
| *abcb1a* | 22141070 | strongHyper | 0.000373 | exon |
| *abcb1a* | 22165829 | strongHypo | 0.0000386 | exon |
| *abcb1a* | 22181973 | strongHyper | 0.000137 | intron |
| *abcb1a* | 22181974 | strongHyper | 0.000138 | intron |
| *abcb1a* | 22182070 | strongHyper | 0.00111 | intron |
| *abcb1a* | 22182101 | strongHyper | 0.00212 | intron |
| *abcb1a* | 22192599 | strongHypo | 0.00392 | intron |
| *abcb1a* | 22222671 | strongHypo | 0.00295 | intron |
| *abcb1a* | 22222675 | strongHypo | 0.00000464 | intron |
| *abcb1a* | 22234557 | strongHyper | 0.00000554 | intron |
| *abcb1a* | 22234706 | strongHyper | 0.0022 | intron |
| *abcb1a* | 22255279 | strongHyper | 0.00128 | intron |
| *abcb1a* | 22279931 | strongHypo | 0.00478 | intron |
| *abcb1a* | 22297010 | strongHyper | 0.0073 | exon |
| *abcb1a* | 22297022 | strongHypo | 0.00601 | exon |
| *abcb1a* | 22304484 | strongHyper | 0.00327 | intron |
| *abcb1a* | 22304577 | strongHyper | 0.00623 | intron |
| *abcb1a* | 22313828 | strongHypo | 0.00286 | intron |
| *abcb1a* | 22320455 | strongHyper | 0.00229 | intron |
| *abcb1a* | 22322568 | strongHypo | 0.00142 | intron |
| *abcb1a* | 22322825 | strongHypo | 0.000596 | intron |
| *abcb1a* | 22397453 | strongHypo | 0.00429 | exon |
| *abcb1a* | 22397481 | strongHypo | 0.000487 | exon |
| *abcb1a* | 22397482 | strongHyper | 0.0000549 | exon |
| *abcb1a* | 22397592 | strongHypo | 0.0018 | exon |
| *abcb1a* | 22141030~22141072 | strongHyper | 0.00000000156 | Exon (exon 25 of 28) |
| *abcb1a* | 22181973~22182103 | strongHyper | 0.000000000029 | Intron (intron 27 of 27) |

### 3.5 Effect of PS on DNA methylation of GC barrier-related genes in the placenta of offspring

To investigate whether the placental GC barrier-related genes P-gp (abcb1a) and FKBP5 gene exhibit differential regulation of gene expression between OPC group and OPS group, their DNA methylation level were further confirmed by Methyltarget sequencing.

A total of 12 placental samples, with six in the OPC group and six in the OPS group, were utilized for validation experiments. The validation results (displayed in Table 4) demonstrate that, P-gp (abcb1a) in the OPS group had 15 differentially methylated CpG sites compared to the OPC group, and three significantly hypermethylated DMR (abcb1a DMR04, abcb1a DMR05 and abcb1a DMR17), FKBP5 in the OPS group had 15 differentially methylated CpG sites, along with two significantly hypermethylated DMR (FKBP5 DMR3 and FKBP5 DMF4).

**Table 4.**
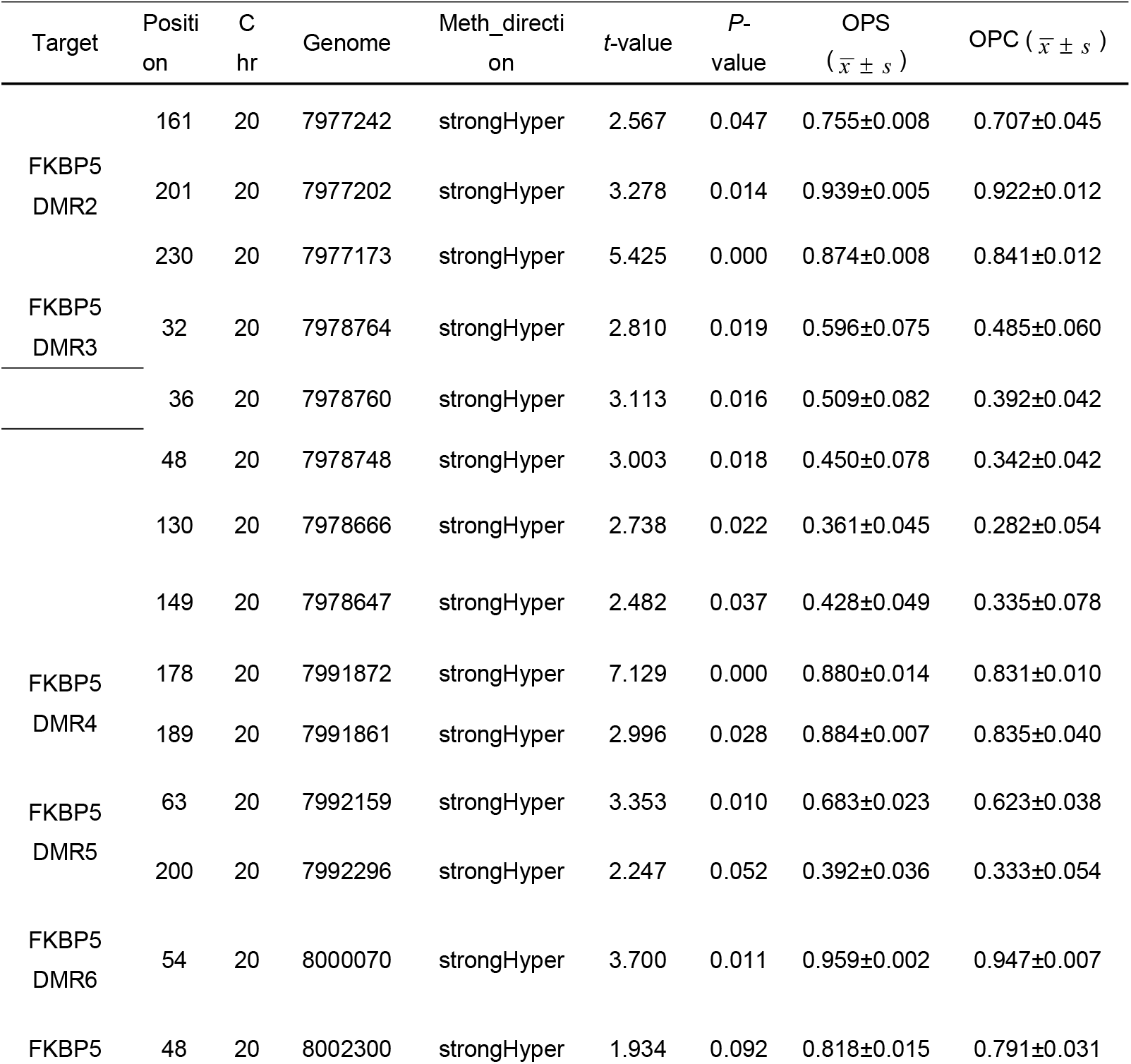
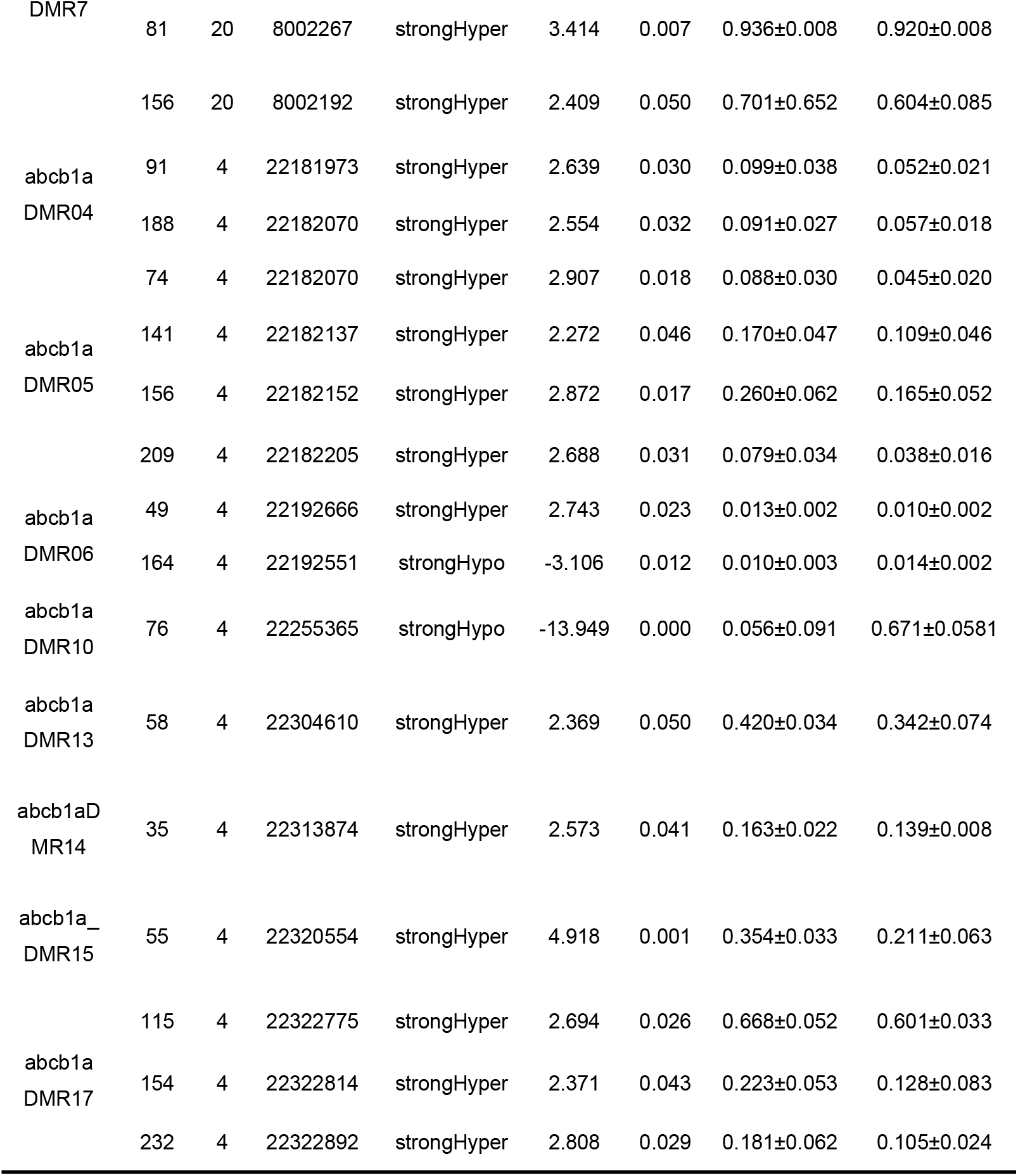
Differences of methylation levels in specific genes CpG site between different groups.

The results also revealed that three identical CpG methylation sites (abcb1a DMR4 chr4:22182070, chr4:22181973, and abcb1a DMR5 chr4:22182070), which were consistent with RRBS results when compared with the OPC group. Then, the CpG sites in each fragment was compared, and nine DMS were found to have the same methylation direction in both sequencing tests. There were 16 DMS in FKBP5, which are different in the two tests, and the methylation directions of three DMS were the same. Furthermore, the FKBP5 gene exhibited significance and hypermethylation in both mean methylation levels and total methylation levels, while the P-gp (abcb1a) gene have significance and hypermethylation in mean methylation levels but not in total methylation levels (Fig.3). Investigation revealed significant differences in methylation levels of P-gp (abcb1a) and FKBP5 as indicated by RRBS and MethylTarget results. Additionally, these methylation sites were found to be distributed within the intron region, suggesting a potential impact of PS on the methylation level of both P-gp (abcb1a) and FKBP5 introns.

**Fig.3.**
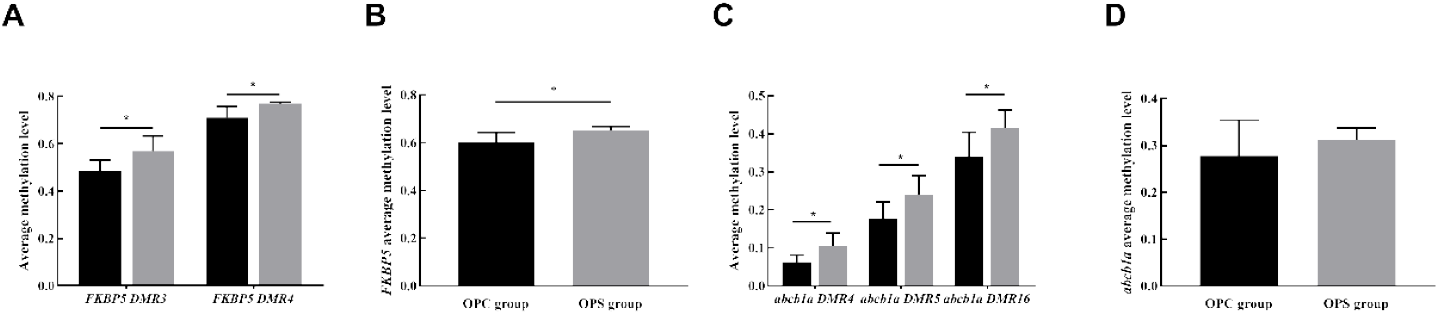
Effects of PS on target regions and total methylation levels of offspring placental GC barrier genes. The figure showed the mean methylation levels and total methylation levels of placental GC barrier-related genes. (A-B) The target mean methylation levels and total methylation levels of FKBP5 were increased by PS on offspring. (C-D) The target means methylation levels of *P-gp (abcb1a)* were increased by PS on offspring but total methylation levels of *P-gp (abcb1a)* were not significant. Data are represented as mean ± SD. Number of animals in each group = 6. *Statistically significant difference compared with the OPC group (*P*<0.05).

### 3.6 Effect of PS on transcription of the placental GC barrier-related genes in offspring

To investigate the mRNA and protein expressions of placental GC barrier-related genes (11β-HSD2, P-gp, NR3C1, and FKBP5), placental samples were collected from both the OPC and OPS groups. Interestingly, a significant reduction in mRNA expression levels of placental *11β-HSD2* (*t*=2.880, *P*=0.016, Fig. 4A) and *P-gp (abcb1b)* (*t*=7.874, *P*=0.000, Fig. 4C) were observed compared to the OPC group. However, there was no statistically significant difference in mRNA levels of *P-gp (abcb1a)* between the OPS and OPC groups (*t*=2.036, *P*=0.081, Fig. 4B). Notably, similar trends were observed at the protein levels of 11β-HSD2 (*t*=5.667, *P*=0.030, Fig. 4F, J), as well as for P-gp (*t*=10.542, *P*=0.009, Fig. 4F, I). Furthermore, FKBP5 mRNA levels were found to be increased in GC-exposed offspring (*t*=-2.428, *P*=0.036, Fig. 4E), while NR3C1 mRNA levels were downregulated (*t*=2.729, *P*=0.032, Fig. 4D). Additionally, the protein expression level of NR3C1 was higher (*t*=-73.099, *P*=0.000, Fig. 4F, I), whereas that of FKBP5 was lower (*t*=12.061, *P*=0.007, Fig. 4F, G), in the OPC group compared to the OPS group. These findings collectively suggest that maternal stress increases the corticosterone concentration in the offspring by weakening the placenta GC barrier function.

**Fig.4.**
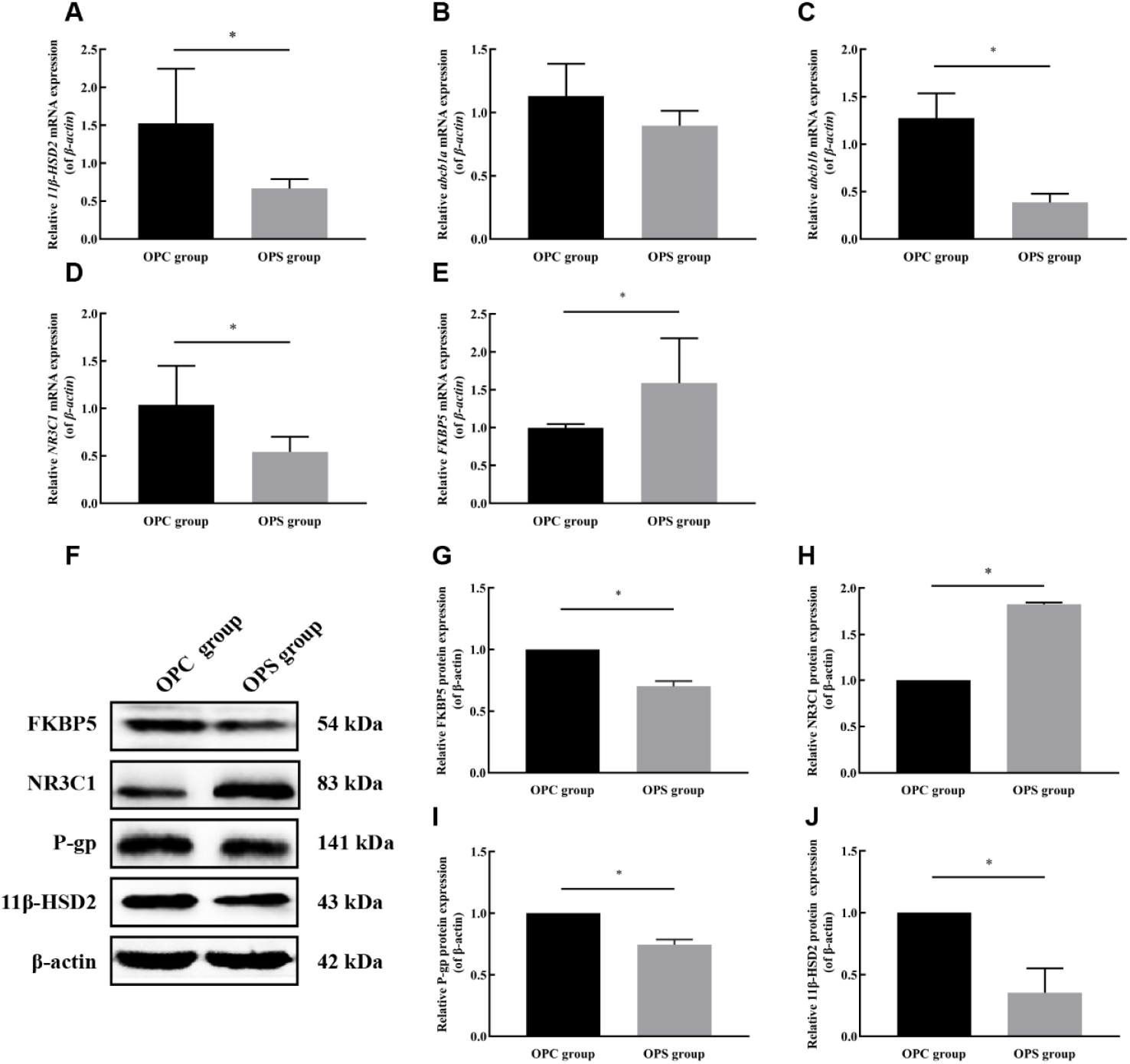
Effects of PS on mRNA and protein expression levels of the offspring placental GC barrier genes. The figure showed that the expressions of 11β-HSD2, P-gp, NR3C1, FKBP5, and β-actin were detected by q-PCR and western blot. (A-D) The mRNA levels of 11β-HSD2, P-gp(abcb1b), and NR3C1 were reduced but FKBP5 expression was increased by PS on offspring. (F-G) The protein levels of 11β-HSD2, P-gp(abcb1b), and FKBP5 were reduced but NR3C1 expression was increased by PS on offspring. Data are represented as mean ± SD. Number of animals in each group = 3. *Statistically significant difference compared with the OPC group (*P*<0.05).

## 4 Discussion

Pregnancy is a sensitive period for maternal and fetal health, with the uterus and placenta critical for fetal development[29]. External stressors can activate the maternal hypothalamic-pituitary-adrenal (HPA) axis, increasing cortisol levels that may cross the placental barrier and affect the fetus[30]. Our study found higher corticosterone levels in prenatally stressed (PS) rats, suggesting HPA axis activation. Excessive corticosterone can potentially cross the placental barrier into the uterus and expose the fetus to elevated levels of this hormone. The placental glucocorticoid (GC) barrier, including 11β-HSD2 and P-gp, protects the fetus from excess cortisol, but PS can downregulate these components, increasing fetal exposure. This can lead to growth retardation and chronic diseases, including neuropsychiatric disorders. These results are consistent with previous studies[19,31,32].

Maternal glucocorticoids regulate the development of the fetal HPA axis by controlling hippocampal GR expression, potentially leading to heightened stress reactivity in adulthood[33]. We verified that using PS produces similar results; in the PS group, mRNA levels of FKBP5 were upregulated, while protein levels were lower compared to the control group. Additionally, mRNA levels of NR3C1 were downregulated and protein levels were higher in the PS group. Although we did not replicate previous findings, our results suggest that an impact on placental GR expression and fetal programming.

Epigenetic changes, influenced by environmental factors, can modify placental function and are linked to PS-induced alterations in genes related to the GC barrier. Our research aligns with prior studies[10,22] found that PS reduced DNMT 3A and 3B expression while increasing DNMT1 suggesting a connection between PS, DNA methylation, and gene expression. MethylTarget analysis identified significant epigenetic differences in the P-gp (abcb1a) and FKBP5 genes, indicating a role for DNA methylation in the placental response to PS.

Research on intrauterine growth restriction (IUGR) implicates epigenetic control of P-gp as pivotal in the placental glucocorticoid (GC) barrier’s function[19]. Our study presents new evidence that P-gp’s role in this barrier is regulated by DNA methylation, which could explain cognitive impairments in offspring due to prenatal stress (PS). Yet, the mechanisms of P-gp’s epigenetic influence need further exploration. Moreover, we found 16 differentially methylated sites (DMS) in the FKBP5 gene, with three showing consistent patterns across tests. Elevated maternal stress was linked to increased methylation of FKBP5, predicting decreased fetal coupling [34]. This finding is consistent with previous research that associates FKBP5 methylation with newborn arousal levels[35,36]. As FKBP5 limits cortisol binding, its increased expression due to methylation changes may modulate cortisol responses.

Increased DNA methylation of the 11β-HSD2 gene, potentially due to maternal prenatal stress, downregulates the placental mRNA and its cortisol-inactivating enzymes[18]. This suggests that prenatal stress modifies the placental barrier’s cortisol regulation, possibly linking to a fetal risk phenotype with reduced coupling[26,37,38]. Despite our study not observing DNA methylation changes in 11β-HSD2 and NR3C1, existing literature indicates that these genes’ methylation correlates with various maternal and neonatal health metrics[23,39]. Additionally, placental NR3C1 methylation has been associated with maternal blood pressure and neonatal outcomes[40]. While our findings on 11β-HSD2 and NR3C1 differ from some reports, they underscore the need for further research on epigenetic regulation within the placenta[36,41]. The study indicates that PS affects the placental GC barrier function through methylation changes in genes like abcb1a and FKBP5, pointing to a broader role for epigenetics in placental function. Ultimately, PS-induced methylation alterations may lead to abnormal glucocorticoid activity and impact offspring growth and development, highlighting the importance of understanding these complex epigenetic influences.

## 5. Conclusions

Our results demonstrate that PS is associated with the methylation and abnormal expression of P-gp (abcb1a) and FKBP5, resulting in the disruption of placental GC barrier integrity and induction of a hypercorticosteroid state in offspring, thereby affecting their growth and development. These suggest that DNA methylation of placental GC barrier genes is the potential early warning target of placental GC barrier opening and susceptibility to multiple diseases.

## Acknowledgments

Thank you to our participants and the Key Laboratory of Environmental Factors and Chronic Disease Control, School of Public Health and Management, Ningxia Medical University for support.

## Data and code availability

The data supporting the findings of this study are available within the article and its supplemental information. The RRBS and MethylTarget sequencing datasets generated in this study can be found in the NCBI SRA database (http://www.ncbi.nlm.nih.gov/sra/) using the access number PRJNA890935 and PRJNA891021.

## CRediT authorship contribution statement

CL, SG, and SM designed and coordinated the study. HL, DY, and RW performed the experiments. HL, YL, and JW analyzed the data. CL and HL wrote the manuscript. All authors contributed to the data discussion and the final manuscript.

